# Molecular mechanism of SARS-CoV-2 cell entry inhibition via TMPRSS2 by Camostat and Nafamostat mesylate

**DOI:** 10.1101/2020.07.21.214098

**Authors:** Tim Hempel, Lluís Raich, Simon Olsson, Nurit P. Azouz, Andrea M. Klingler, Marc E. Rothenberg, Frank Noé

**Author notes:** Equal contribution (Lluís Raich, Simon Olsson).

## Abstract

The entry of the coronavirus SARS-CoV-2 into human cells can be inhibited by the approved drugs camostat and nafamostat. Here we elucidate the molecular mechanism of these drugs by combining experiments and simulations. *In vitro* assays confirm the hypothesis that both drugs act by inhibiting the human protein TMPRSS2. As no experimental structure is available, we provide a model of the TMPRSS2 equilibrium structure and its fluctuations by relaxing an initial homology structure with extensive 280 microseconds of all-atom molecular dynamics (MD) and Markov modeling. We describe the binding mode of both drugs with TMPRSS2 in a Michaelis complex (MC) state preceding the formation of a long-lived covalent inhibitory state. We find that nafamostat to has a higher MC population, which in turn leads to the more frequent formation of the covalent complex and thus higher inhibition efficacy, as confirmed *in vitro* and consistent with previous virus cell entry assays. Our TMPRSS2-drug structures are made public to guide the design of more potent and specific inhibitors.

## 1 Introduction

In December 2019 several cases of unusual and severe pneumonia were reported in the city of Wuhan, China. These cases were traced back to a new coronavirus, SARS-CoV-2 (Severe acute respiratory syndrome coronavirus 2); the disease is called COVID-19 [1]. As of July 1st 2020 there are over 10 million confirmed COVID-19 cases and more than 500,000 deaths [2], with both numbers likely to be severe underestimates. Given estimates of the infection mortality rate of 0.4 to 1.4 % [3–5] the virus has the potential to kill tens of millions of people unless efficient vaccines or drugs are available.

As other coronaviruses [6–9], SARS-CoV-2, exploits host proteins to initiate cell-entry, in particular TMPRSS2 and ACE2, two membrane-bound proteins expressed in the upper respiratory tract. TMPRSS2 contains an extracellular trypsin-like serine-protease domain that is thought to be essential for activation of the spike (S) protein on the surface of SARS-CoV-2 viral particles [10] (Fig. 1a). Together with its activation, the binding of S-protein to ACE2 [11], triggers the cell-entry process [12]. TMPRSS2 is also required by other viruses, including several strands of influenza [13, 14]. Moreover, epidemiological data of prostate cancer patients undergoing androgen-deprivation therapies, which lowers TMPRSS2 levels, indicate a lower risk of contracting the SARS-CoV-2 infection [15]. TMPRSS2 knockout mice have no severe phenotype [16], indicating that inhibiting TMPRSS2 function may not lead to adverse side-effects. Consequently, TMPRSS2 is a promising therapeutic targets to stifle or block coronavirus infection, while simultaneously maintaining a low risk of drug resistance development.

**Figure 1:**
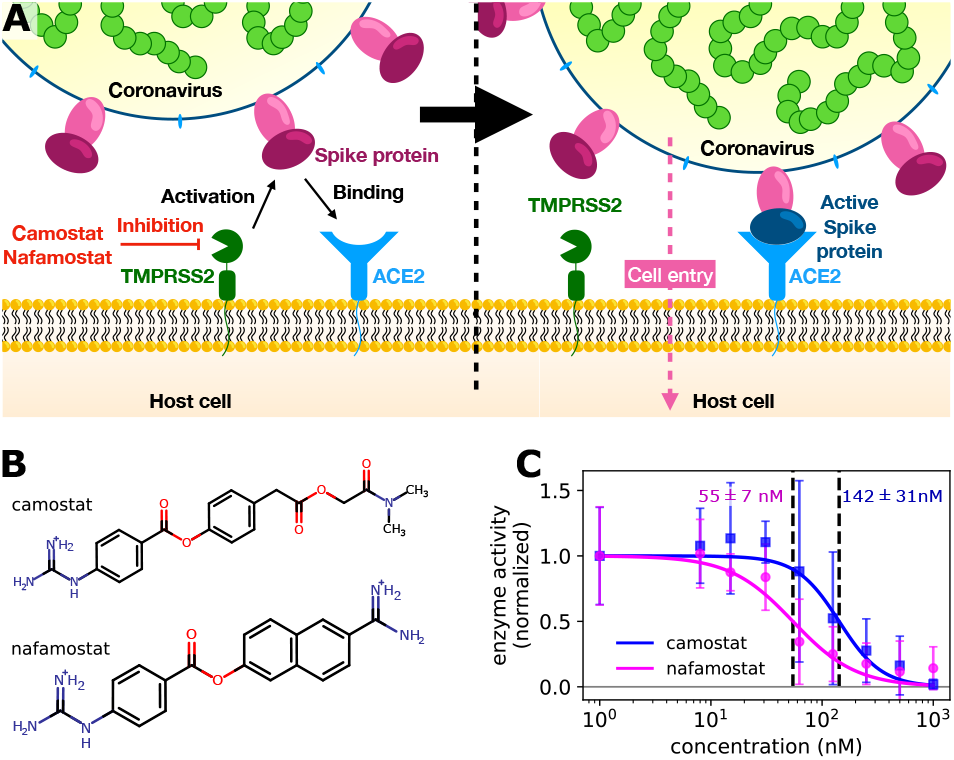
**A** overview of viral entry. **B** structures of tested drugs in native protonation state. **C** Dose response behavior of TMPRSS2 inhibition by camostat and nafamostat with computation of IC50 (normalized, background subtracted).

Here, we study the structural basis and molecular mechanism of TMPRSS2 inhibition by camostat and nafamostat. Both guanidinobenzoate-containing drugs are approved in Japan and have been demonstrated to inhibit SARS-CoV-2 cell-entry [10, 17, 18]. We report experimental measurements demonstrating that these drugs inhibit TMPRSS2 activity by using our recently established cell-based assays [19], consistent with *in vitro* enzymatic TMPRSS2 activity assays [20].

While TMPRSS2 is an excellent drug target against SARS-CoV-2 and other viruses, we are, as yet, lacking an experimental structure. We here go beyond the previous dependency on homology models by an extensive 280 microseconds of high-throughput all-atom molecular dynamics (MD) simulations and Markov modeling. This approach provides an ensemble of equilibrium structures of the protein-drug complex and also drug binding kinetics. Nafamostat and camostat are both covalent inhibitors with an identical covalent complex, but the greater inhibitory activity of nafamostat can be explained by a greater stability of its Michaelis complex preceding the covalent complex. These findings, combined with the simulation structures that we make publicly available, provide a keystone for developing more potent and specific TMPRSS2 inhibitors.

## 2 Results

### Camostat and Nafamostat inhibit the catalytic activity of TMPRSS2

First we confirm the hypothesis that camostat and nafamostat inhibit cell entry of SARS-CoV-2 and other coronaviruses by inhibiting the TMPRSS2 protein. To this end, we use our recently reported assay [19] of full-length TMPRSS2 activity on the surface of live cells with both inhibitors (Fig. 1B). Briefly, we transfected the human cell-line HEK-293T with TMPRSS2. We then measured the protease activity of the transfected cells using the fluorogenic peptide substrate BOC-QAR-AMC, following incubation of the cell with increasing inhibitor concentrations. Peptide-digestion induced a minimal increase in fluorescent signal in control cells without exogenous TMPRSS2 expression (unnormalized mean enzyme activity = 2.4), while TMPRSS2 over-expression resulted in a much faster peptide digestion (normalized mean enzyme activity = 12.8). Therefore, our assay is mostly specific for TMPRSS2 [19]. Significantly lower enzyme activity at higher drug concentrations can thus be attributed to TMPRSS2 inhibition.

For both camostat and nafamostat, we see a clear dose-dependent inhibition and estimate their respective IC50 values to 142 ± 31 nM and 55 ± 7 (Fig. 1C). Our results are consistent with the finding that both drugs inhibit cell entry of SARS-CoV-2 and other coronaviruses, and that nafamostat is the more potent inhibitor [17, 18, 20].

### Equilibrium structures of TMPRSS2 in complex with Camostat and Nafamostat

We now set off to investigate the molecular mechanism of TMPRSS2 inhibition by camostat and nafamostat. While no TMPRSS2 crystal structure is available to date, it has been shown that all-atom MD simulations can reliably model the equilibrium structures of proteins when (i) a reasonable model is available as starting structure, and (ii) simulations sample extensively, such that deficiencies of the starting structure can be overcome [22–26].

Here, we initialize our simulations with recent homology models of the TMPRSS2 protease domain and with camostat/nafamostat docked to them [27]. Trypsins adopt a common fold and share an active-site charge relay system whose structural requirements for catalytic activity are well understood [28], and we selected protein consistent with these structural requirements. In particular, we focused on systems with Asp435 deprotonated and His296 in a neutral form (N_*δ*_ protonated), as well as on the interactions of a charged lysine nearby the catalytic Asp345 (Figs. S1,S2).

In order to overcome artifacts of the initial structural model and simulate the equilibrium ensemble of the TMPRSS2-drug complexes, we collected a total of 100 *μs* of simulation data for TMPRSS2-camostat and 180 *μs* for TMPRSS2-nafamostat. Both simulations sample various drug poses and multiple association / dissociation events. Using Markov modeling [29–33] we elucidate the structures of the long-lived (metastable) states and characterize, protein-drug binding kinetics and thermodynamics.

We find TMPRSS2 has flexible loops around the binding site but to maintain stable structural features shared by other trypsin-like proteases (Fig. 2A). These enzymes cleave peptide-like bonds in two catalytic steps, assisted by a conserved catalytic triad (Asp345, His296, and Ser441 in TMPRSS2). The first step involves the formation of a covalent acyl-enzyme intermediate between the substrate and Ser441 [28]. During this step His296 serves as a general base to deprotonate the nucleophilic Ser441, and subsequently as a general acid to protonate the leaving group of the substrate. The second step involves the hydrolysis of the acyl-enzyme intermediate, releasing the cleaved substrate and restoring the active form of the enzyme.

Along these two steps, the so called “oxyanion hole”, formed by the backbone NHs of Gly439 and Ser441, helps to activate and stabilize the carbonyl of the scissile bond. Another important structural feature is the S1 pocket, which contains a well conserved aspartate (Asp435) that is essential for substrate binding and recognition. At the opposite site of the S1 pocket, a loop containing a hydrophobic patch delimits the binding region of substrates within enzymatic active site. All these structural elements, known to play crucial roles in the function of serine proteases [28], are generally stable and preserved in our equilibrium structures.

**Figure 2:**
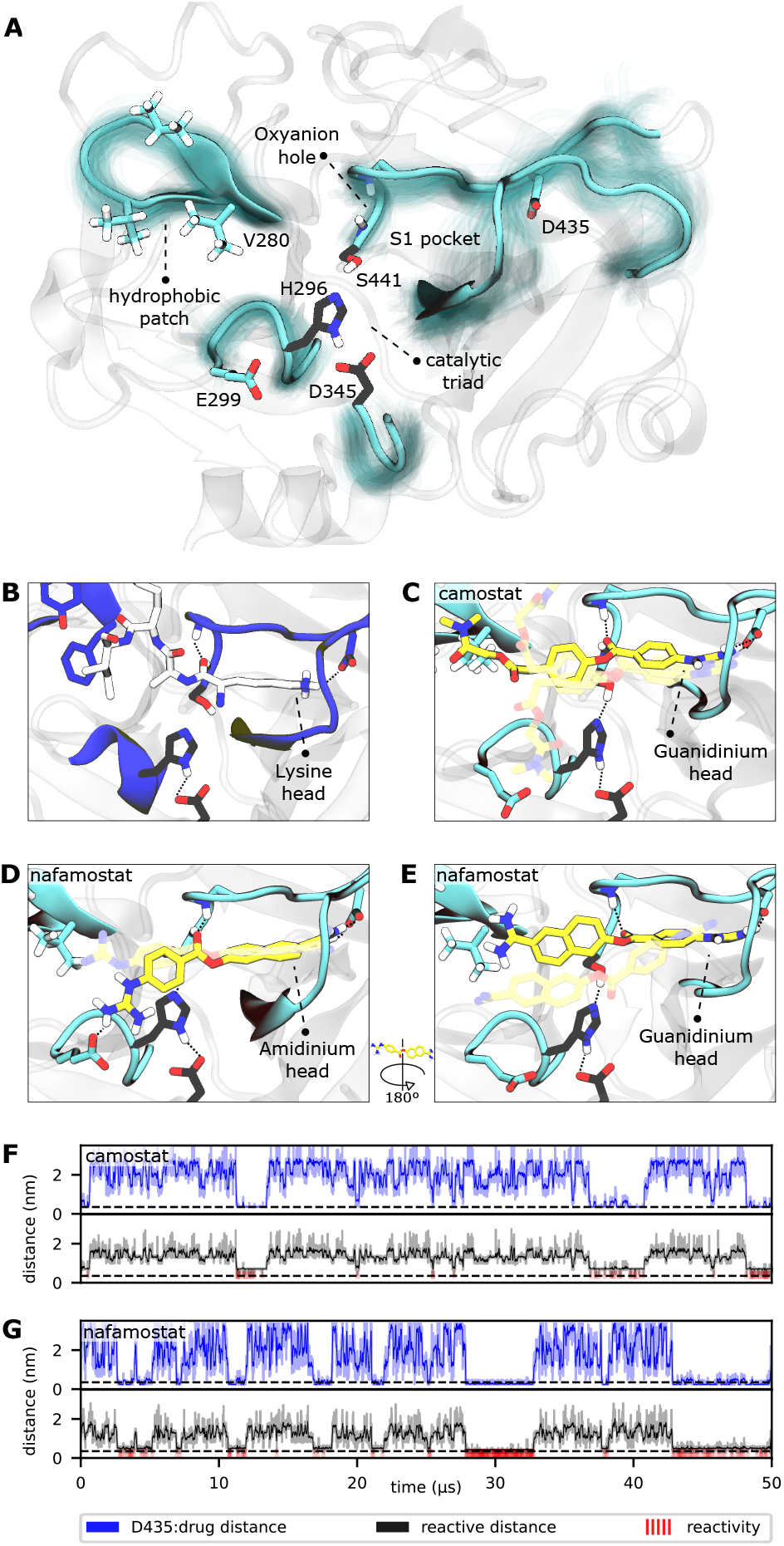
TMPRSS2 structure and Michaelis complex with camostat and nafamostat. **A**: Overview of catalytic domain of TMPRSS2 simulation structures and detailed view of active site, catalyic triad shown in black. **B**: substrate processing shown at example of trypsin (PDB ID 4Y0Y [21]). **C**: camostat associated to TMPRSS2; guanidinium head is interacting with D435 in S1 pocket. **D+E**: nafamostat associated to TMPRSS2 with aminidinium (**D**) or guanidinium (**E**) heads interacting with D435. **F+G**: Markov model simulations of minimal camostat (**F**) and nafamostat (**G**) distance to D435 (S1 pocket binding, blue) and reactivity coordinate (mean of drug ester carbon to catalytic serine oxygen and catalytic serine hydrogen to catalytic histidine nitrogen). Reactivity, i.e. when both reactive distances are within range, is indicated with red markers.

### Structural basis of TMPRSS2-inhibition by Camostat and Nafamostat

Drugs with a guanidinobenzoate moiety can inhibit trypsins by mimicking their natural substrates (Fig. 2B). Indeed, the ester group, resembling a peptide bond, can react with the catalytic serine with rates that are orders of magnitude faster [34], forming the acyl-enzyme intermediate. In contrast to peptide catalysis, the drug’s guanidinobenzoyl group stays covalently linked to the catalytic serine with a small off-rate, rendering it an effective chemical inhibitor [35].

In our MD simulations, we sample different conformations of the complex formed by the enzyme and each of the drugs, and we can thus elucidate how they bind and how specificity (or lack thereof) arises before the catalytic step. They primarily mimic interactions made between trypsins and their substrates, with lysine heads interacting with a conserved asparatate in the S1 pocket (Asp435, Fig. 2B).

A fraction of the bound-state structures resembles a reactive Michaelis complex (MC), with the guanidinium head interacting with Asp435 (for both camostat and nafamostat) as well as a flipped orientation where the amidinium head of nafamostat, also charged, docks into the S1 pocket and interacts with Asp435 (Fig. 2C,D,E). Interestingly, in this inverted orientation of nafamostat, the guanidinium head mainly interacts with Glu299, with the drug reactive center slightly displaced from the oxyanion hole, while the other orientation keeps the amidinium head mainly nearby Val280, with the ester center well positioned for the reaction. This observation is in agreement with several crystal structures of acylenzyme intermediates between different trypsins and guanidinobenzoyl molecules bound to the S1 pocket (e.g. PDBs 2AH4 [36], 3DFL [37], 1GBT [38]). There are also “inverse substrates” known to react with rates comparable to the ones of normal esters, suggesting that the inverted nafamostat orientation may also be reactive [28].

### Kinetic mechanism of TMPRSS2-inhibition by Camostat and Nafamostat

Finally, we investigate the molecular basis for the greater inhibition by nafamostat and formulate starting points for designing new and more efficient covalent TMPRSS2 inhibitors following these leads.

To illustrate the reversible binding of camostat and nafamostat to TMPRSS2 we used our Markov models to simulate long-time scale trajectories of 50 *μs* (Fig. 2F,G). We see a clear correlation between the formation of a tight interaction to Asp435 in the S1 pocket to the inhibitor, and the ester group of the inhibitors into close contact with the catalytic serine, thereby forming a reactive complex. In other words, the binding of reactive drugs in the S1 pocket favors the interactions that lead to a catalytically competent MC.

We estimate the dissociation constant for the MC, i.e. the ratio of dissociated state and MC population, to be 5.95 mm (5.64, 78.3) for camostat and 6.07 mm (6.033, 28.6) for nafamostat (67% confidence intervals). Comparing to an experimental estimate of the overall dissociation constant of *K_i_* = 11.5 μm for nafamostat to bovine pancreatic trypsin [35], this indicates that the major source of inhibition is not the non-covalent MC, but rather the longer-lived covalent acyl-enzyme complex. However, as camostat and nafamostat yield identical acyl-enzyme complexes, their differences can only arise from either (1) the formation or population of their MCs, or (2) differences in the catalytic rate *k*_cat_ of acylation.

Interestingly, we observe a three-fold higher population of the of the MC with nafamostat compared to camostat, as well as a significantly higher on-rate (Fig. 3). A simple three-state kinetic model of dissociated state, MC and covalent complex shows that the overall association constant (*K_a_*, ratio of inhibited versus apo protein states) directly scales with the association constant of the MC (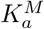, ratio of MC versus dissociated states) by a constant factor (Methods):

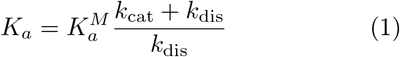

**Figure 3:**
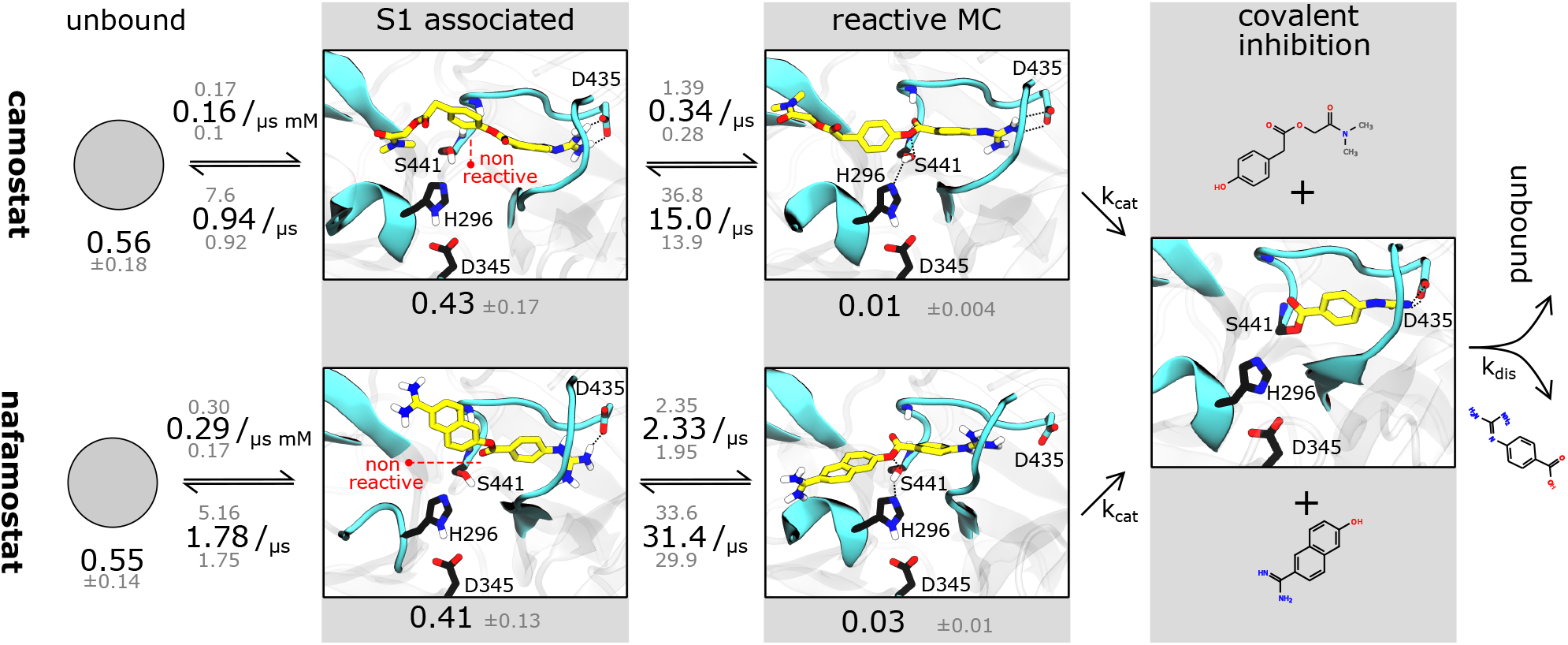
Binding rate model of camostat (top) and nafamostat (bottom). Inhibition process is depicted from left to right. Single representative structures are shown for the sake of readability, we however note that states show flexibility. Rates and populations predicted by our model are annotated to reaction arrows and states. Covalent inhibition by nafamostat binding group depicted at example of prostatin (PDB 3DFL [37]).

Simply speaking, this indicates that nafamostat is a better inhibitor because it is more often found the reactive MC state, and is therefore more likely to be attacked by the catalytic serine oxygen and enter the long-lived acyl-enzyme inhibitor complex.

Moreover, we note that the *k*_cat_ of acylation of these drugs may depend on their leaving group pKa’s. Indeed, leaving groups with a low pKa will require less assistance from acid catalysis and will be easily displaced by the nucleophilic serine, favoring the formation of the acyl-enzyme intermediate. We expect the leaving group of nafamostat to have a lower pKa than the one of camostat, following the values of similar molecules such as naphtol (9.57 [39]) and 4-methylphenol (10.26 [40]), respectively. This suggests that the *k*_cat_ of acylation will be also slightly faster for nafamostat, further contributing to its superior inhibition of TMPRSS2.

## 3 Discussion

Camostat and nafamostat are promising drug candidates for a COVID-19 drug treatment strategy. Here we have combined cell-based assays, extensive molecular simulations, and Markov modeling to unravel the molecular action principle of these drugs and provide data that may help to improve them further.

Our binding assays provide evidence that both inhibitors directly act on TMPRSS2 and that nafamostat is more potent compared to camostat, and this qualitative difference is in agreement with complementary *in vitro* assays on purified protein [20] cell-entry assays [17, 18]. We note that the absolute IC50s values differ between these three assay types, reflecting differences in experimental conditions and which function is being inhibited and measured.

While no crystallographic structure of TMPRSS2 is available, we provide extensive 280 microseconds of all-atom MD simulations starting from a homology model that generate stable equilibrium structure ensembles of the two protein-drug complexes. These simulations sample multiple association / dissociation events and various drug poses in the protein active site. Our analyses show that the non-covalent complexes of camostat and nafamostat with TMPRSS2 are relatively short-lived, suggesting that the main inhibitory effect is due to the formation of the long-lived covalent acyl-enzyme complex between the drug’s guanidinobenzoyl moiety and the catalytic serine of TMPRSS2.

Although the MC state is not the main cause of inhibition, its population directly translates into the potency of the inhibitor as higher MC population corresponds to a higher catalytic rate and therefore a higher overall population the protein spends locked down in the covalent inhibitory complex. Consistently with the higher potency of nafamostat, it is found to have a threefold more stable MC compared to camostat. A second contribution may be the pKa of drug leaving groups, affecting the rate of enzyme acylation.

Our detailed models of the thermodynamic and kinetics of inhibitor binding highlight the bound state’s heterogeneity, with both drugs adopting multiple distinct poses. We note the importance of residue Asp435 in the conserved S1-pocket, which stabilizes the MC state and helps to orient the reactive molecules in a conformation that is suited for catalysis. Nafamostat has two groups that can potentially bind into the S1 pocket, whereas camostat has only one. However, we find that the population of S1 associated states are similar between camostat and nafamostat, suggesting that non-covalent inhibition is likely a minor contribution to the overall inhibition of TMPRSS2.

We conclude that the design of future TMPRSS2 inhibitors with increased potency and specificity should incorporate the following points

First, stabilizing the non-covalent complex with the TMPRSS2 active site is beneficial for both, covalent and non-covalent inhibitors. As S1 pocket binding is a major contribution to the stability of the non-covalent complex, effective drugs may contain hydrogen bond donors and positively charged moieties that could interact principally with Asp435, but also with different backbone carbonyls of the loops that compose the cavity (e.g. from Trp461 to Gly464).

Second, for covalent inhibitors, we must consider that the catalytic serine is at a distance of around 1.3 nm from Asp435. Thus the reactive center of an effective drug and its S1-interacting moieties should be within that distance. We further suggest that a drug should be size-compatible to the hydrophobic patch on the S1 distal site (Fig. 2A). We speculate that drugs with a large end to end distance and high rigidity may not fit well in the described TMPRSS2 scaffold, and in particular, might be significantly less reactive.

Third, optimizing the pKa of the drug’s leaving group might be beneficial for improving covalent TMPRSS2 inhibitors. The first step of the reaction would be faster, and the acetyl-enzyme intermediate would accumulate. We note that the deacetylation off-rate must be very low, ideally on the order of magnitude of guanidinobenzoate moiety containing drugs.

Finally, we make our simulated equilibrium structures of TMPRSS2 in complex with camostat and nafamostat available, hoping they will be useful to guide future drug discovery efforts.

## Acknowledgements

We acknowledge financial support from Deutsche Forschungsgemeinschaft DFG (SFB/TRR 186, Project A12), the European Commission (ERC CoG 772230 “ScaleCell”), the Berlin Mathematics center MATH+ (AA1-6) and the federal ministry of education and research BMBF (BIFOLD). We are grateful for in-depth discussions with Stefan Pöhlmann (DPZ Göttingen), Katarina Elez, Robin Winter, Tuan Le, Moritz Hoffmann (FU Berlin) and the members of the JEDI COVID-19 grand challenge.

## Software and data availability

Structural ensembles of camostat and nafamostat binding poses are published online at https://github.com/noegroup/tmprss2_structures.

## Materials and Methods

### TMPRSS2 activity assays

TMPRSS2 activity assay was described previously [19]. Briefly, we transfected HEK-293T with a PLX304 plasmid containing the open reading frame (ORF) sequence of TMPRSS2 which encodes for the full length protein (492 amino acids). Control experiments are conducted with PLX304 plasmids.

Eighteen hours later, we replaced the media to either PBS alone or PBS in the presence of varying concentrations of candidate inhibitors camostat and nafamostat. Fifteen minutes later, we added the fluorogenic substrate BOC-QAR-AMC to the wells to induce a measurable signal of enzyme activity. We measured the fluorescent signal immediately after adding the substrate, in 15 minutes intervals for a total time of 150 minutes [19]. A baseline proteolytic activity of control cells was measured; we hypothesize that this is because of proteolytic cleavage of the substrate by endogenous transmembrane proteases. However, the TMPRSS2 overexpression cells have significantly increased proteolytic activity compared with control cells [19].

To validate the exogenous expression of TMPRSS2, we performed western-blot analysis of cell lysates from TMPRSS2 overexpressing cells and control cells. A 60kDa band was observed in TMPRSS2 overexpressing cells but not in control cells, which is the expected molecular weight of TMPRSS2 protein after post transcriptional modifications, indicating that the target protein has been successfully expressed.

### IC50 estimation

We used a generalized log-logistic dose-response model

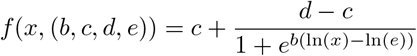

with the concentration *x*, *c* and *d* representing the lower and upper limits, *b* steepness of the curve, and *e* to estimate IC50 values [41].

Upper and lower limits were set to the means computed from control experiments with no drug (upper limit) and PLX plasmid (no TMPRSS2; background noise). We used scipy’s [42] curve fitting algorithms to extract the IC50 with error estimates.

### Molecular dynamics simulations

MD simulations were run with OpenMM 7.4.0 [43] using the CHARMM 36 force field (2019 version) [44]. Camostat and nafamostat structures were taken from PubChem [45] with PubChem CIDs 4413 (nafamostat) and 2536 (camostat), respectively, and modeled with the CHARMM general force field (CGenFF v. 4.3) [46]. We generated our MD setups with CharmmGUI [47]. We initiate a simulation box of side length 7.5 nm with a NaCl ion concentration of 0.1 mol/l at neutral charge and the TIP3P water model [48]. The setups contain 12038 (camostat) and 12039 (nafamostat) water molecules.

We run simulations in the NPT ensemble and keep the temperature at 310 K (physiological temperature) and the pressure at 1 bar. We use a Langevin integrator with 5 fs integration step and heavy hydrogen approximation (4 amu). PME electrostatics, rigid water molecules, and a 1 nm cutoff for non-bonded interactions are used. Simulation times vary between 100 and 500 ns and accumulate to 100 *μs* (camostat) and 180 *μs* (nafamostat), respectively. Structures were visualized using VMD [49].

Due to the lack of a crystal structure for TMPRSS2, MD simulations were seeded from a homology model. It is taken from Ref. [27], model 3W94 is chosen based on precursive MD analyses that showed that 3W94 has the most stable catalytic triad configuration (Figs. S1,S2). The construct includes amino acids 256 to 491 of the full sequence, corresponding to the catalytic chain except for a C-terminal Glycine missing due to homology modeling against a shorter sequence. MD simulations are seeded as follows: Equilibrated docking poses (highest scorers of Ref. [27]) of the ligand were generated in a precursive run using another homology model. We note that the used camostat docking pose resembles the one described by [50]. This data set was equilibrated with local energy minimization, 100 ps simulations with 2 fs time steps in NVT and NPT ensemble subsequently. Frames are selected based on a preliminary metastability analysis, protein conformation is constraint to 3W94 homology model using a constraint force minimizing minRMSD. Production run MD simulations are started from these poses, i.e. from the same protein configuration and with 77 (nafamostat) and 60 (camostat) ligand docking poses, respectively. To ensure convergence of sampling statistics, we ran multiple adaptive runs of simulations, seeding new simulations with coordinates associated with sparsely sampled states.

### Markov modeling

We model the binding and unbinding rates in a two step procedure using Markov state type models [29–32, 51–53]. First, we describe drug unbound and associated states using a hidden Markov model (HMM) [54]. Second, we define a reactive state by using distance cutoffs.

In detail, in the first stage we define distance features between drug guanidinium group and TMPRSS2 Asp435 (minimal distance), drug amidinium group and TMPRSS2 Asp435 (minimal distance, nafamostat only). We further use a binary “reactive” distance feature defined by drug ester carbon to catalytic Ser441-OG, and catalytic serine (HG) to catalytic histidine (NE2) and a threshold of 0.35 nm. If both last mentioned distances are below the threshold, both nucleophilic attack of the serine-OG to the drug ester group and proton transfer from serine to histidin are possible, thus defining the reactive state.

We discretize this space into 240 (camostat) and 490 (nafamostat) states using regular spatial clustering and use an HMM at lag time 5 ns with 5 (camostat) or 8 (nafamostat) hidden states. Nafamostat yields two S1 associated states encoding for both binding directions, camostat a single one, that are defined by being at salt bridge distance to Asp435. We note no significant correlation between the hidden states and the reactive state, i.e. reactivity is not metastable. The described HMMs are used to generate the time series presented in Fig. 2. Reactivity according to this basic HMM does not necessitate S1 pocket binding.

In the second stage, we bisect the HMM binding poses into reactive and non-reactive by combining HMM Viterbi paths [55] and the reactive state trajectories to one single discrete trajectory consisting of 3 states. We define the S1 associated states by filtering the Viterbi paths of the HMM according to S1-association. We use the reactivity trajectories to further bisect the S1 associated state into reactive and non-reactive states, yielding a three state discretization of the drug binding mode. Note that the S1-reactive state is a subset of the reactive state in the plain HMM model (stage 1).

We estimate a maximum likelihood Markov state model (MSM) from the stage 2 trajectories. We report the stationary probability vector as well as transition rates. The latter are approximated using the matrix logarithm approximation of scipy [42] to compute the transition rate matrix *R* from the transition probability matrix T using the definition *T* = exp(*Rτ*) with the lag time *τ*. We found that all reported quantities are converged with respect to the lag time above *τ* = 500 ns which was thus chosen as the model lag time. Errors are estimated by bootstrapping validation using a random subset of 80% of the data (without replacement). All MSM/HMM analyses were conducted using the PyEMMA 2 software [56].

### Kinetic model

Simplifying the binding kinetics into a three-state model describing the binding to / dissociation from the Michaelis complex (ligand concentration c and rates *k*_on_, *k*_off_), catalytic rate of entering the covalent complex (*k*_cat_) and dissociation to the apo state (*k*_dis_), the kinetics are described by the rate matrix:

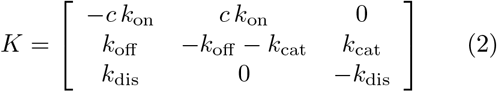

with the (unnormalized) equilibrium distribution

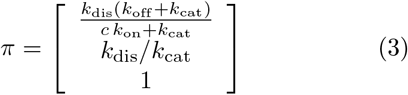

The overall dissociation constant is then:

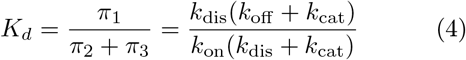

The non-covalent dissociation constant of the Michaelis complex:

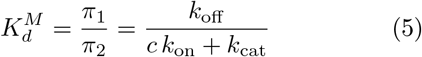

The dissociation constant scales as:

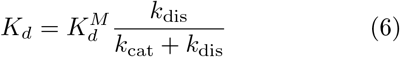

And thus the association constant scales with the stability of the Michaelis complex by a constant factor given by the rates of chemical catalysis and dissociation:

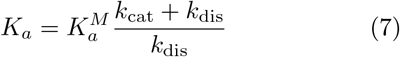

---

**Figure S1:**
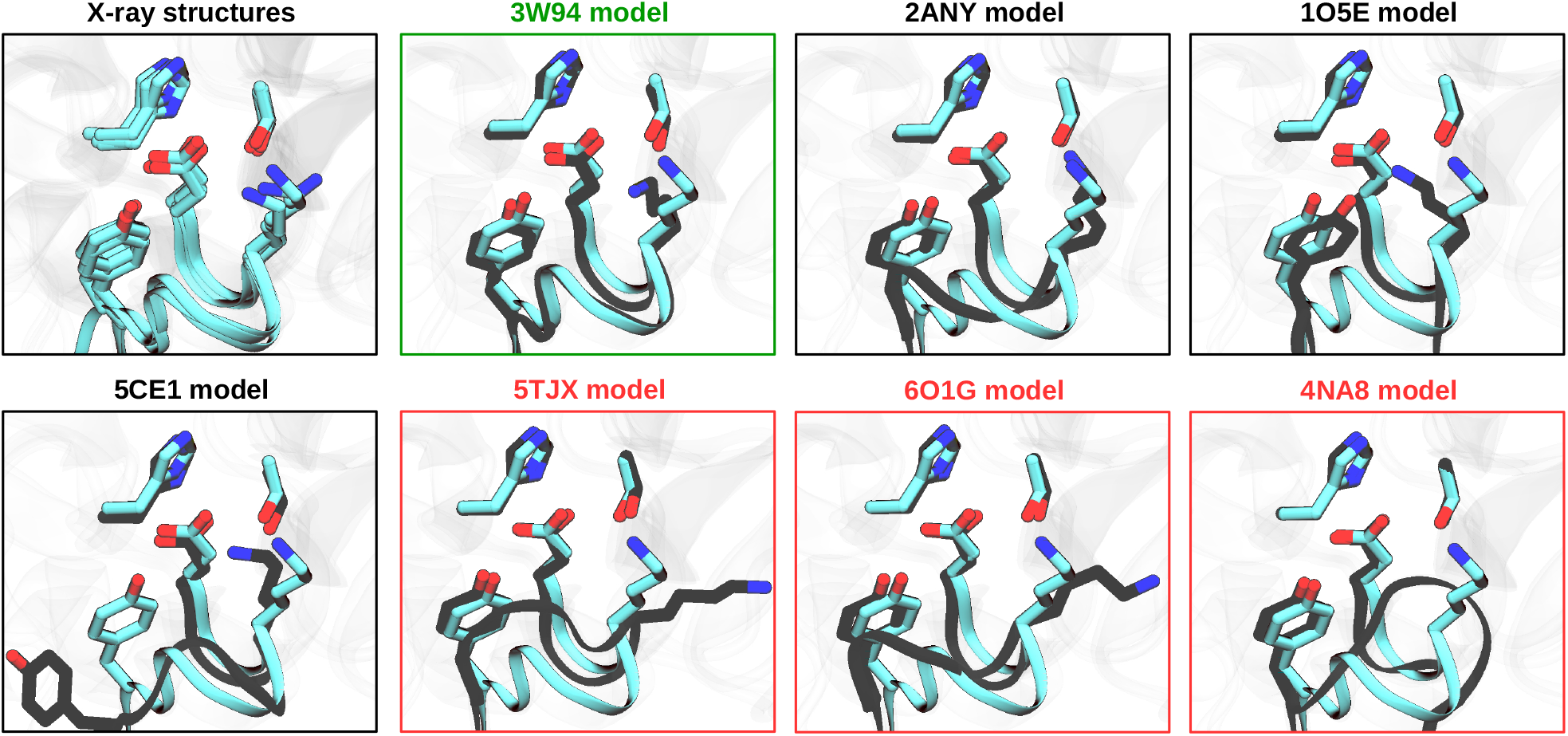
Selection of homology model from Ref. 27 by specific interactions around the catalytic triad. Comparison of TMPRSS2 models (black) and four serine protease structures that contain a lysine residue next to the catalytic aspartate (cyan, PDBs 1EKB, 1FUJ, 3W94 and 4DGJ). Note that the four crystal structures show a very conserved and rigid environment around the catalytic aspartate, with just few fluctuations of the lysine head (K99 in 1EKB). The structural models of TMPRSS2, instead, show a wide variability of conformations of both the backbone and sidechains, with 3W94 being the most conservative model.

**Figure S2:**
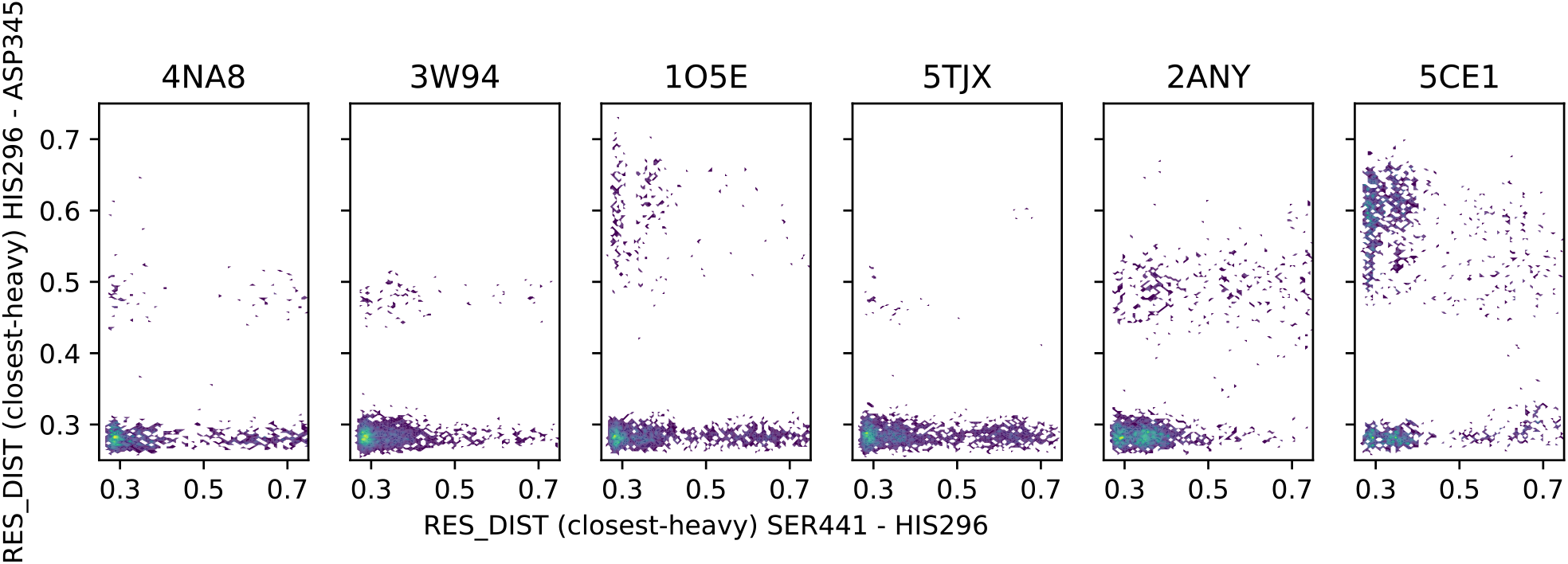
Relevant distances of the catalytic triad for different homology models from Ref. [1] as computed from roughly 30*μs* of MD data for the drug free protein.

## Notes

### Competing Interest Statement

The authors have declared no competing interest.

